# Neural subspaces of imagined movements in parietal cortex remain stable over several years in humans

**DOI:** 10.1101/2023.07.05.547767

**Authors:** L Bashford, I Rosenthal, S Kellis, D Bjånes, K Pejsa, BW Brunton, RA Andersen

## Abstract

A crucial goal in brain-machine interfacing is long-term stability of neural decoding performance, ideally without regular retraining. Here we demonstrate stable neural decoding over several years in two human participants, achieved by latent subspace alignment of multi-unit intracortical recordings in posterior parietal cortex. These results can be practically applied to significantly expand the longevity and generalizability of future movement decoding devices.

## Main Text

Brain-machine interfaces (BMIs) decode neural activity to reproduce the user’s intention and assist individuals with physical and neurological disabilities. In motor BMIs, the user commonly imagines or attempts to make a movement, and the corresponding recorded neural activity is decoded to guide movement in the intended direction, either on a computer or a prosthetic ^1,2^. BMIs can use neural signals acquired at different spatial and temporal resolution, but these have tradeoffs in performance and stability. Whereas single- or multi-unit recordings provide the highest information content, these recordings suffer from non-stationarity – different individual neurons are recorded from day to day or even morning to afternoon^3,4^. This variation is caused by several factors, including movement of the electrodes, changes in the electrode-tissue interface, and degradation of the electrodes. Thus, as the neural features used to train the decoder change, the performance of the BMI degrades over time. As BMIs are implanted for increasingly long durations ^5,6^ the longitudinal stability of intracortically recorded neural activity is a central challenge to the practical utility of BMI devices. Currently, long-term use of BMI devices is only possible when users perform frequent retraining, often several times in a single day, to maintain desired performance. In addition to being time consuming, frequent retraining may not be possible in some use cases, for example in degenerative diseases (e.g., ALS/MND – amyotrophic lateral sclerosis/motor neuron disease) where the loss of function over time may eventually prevent the performance of training tasks.

Although different neurons from the same population are being recorded, the lower-dimensional subspaces of the neural dynamics may remain relatively stable ^7,8^; we investigate this intriguing possibility in the context of BMI decoding. Alternative neural signals such as the local field potential have been observed to be more stable over time ^9,10^; however, the tradeoff is a reduction in information content compared to single unit activity, which ultimately limits decoding performance. Therefore, the most promising solution currently being investigated is to use ‘latent signals’ for BMIs. Latent signals are derived from low-dimensional subspaces of the original high-dimensional single- or multi-unit (MUA) neural activity, and they have been shown to preserve information content while minimizing non-stationarity ^11–15^. Most stable latent spaces have so far been identified and validated primarily in longitudinal recordings of non-human primate primary sensorimotor cortices. In this paper, we investigate the potential of these latent signals for BMI control in two human participants, for whom neural signals were recorded in higher order cortical areas over several years ^16,17^. Specifically, we demonstrate that the neural subspace of imagined reaches in a center-out task remained remarkably stable in posterior parietal cortex.

Data were collected on 143 and 73 unique days, aggregated over a total period of 1106 and 871 days, for participant 1 ^1^ and 2 ^18^, respectively. Participant 1 attempted reaches in 4 directions while MUA was recorded from Brodmann Area 5 (BA5) and the Anterior Intraparietal Area (AIP). Participant 2 attempted reaches in 8 directions while MUA was recorded from the junction of the postcentral and intraparietal sulcus (PC-IP) (Fig. 1A). We use only the ‘training’ trials for longitudinal analysis, without any decoder present, to ensure the data were directly comparable from day-to-day ^19^. During these trials, participants imagined moving their arm to follow the movement of an on-screen cursor. To process the neural data, we adapted methods established in non-human primates ^12^. First a latent signal for each day on which the experiment occurred is calculated by performing Principal Component Analysis (PCA) ^7^ on all trials that day. The latent signal is then aligned for all pairs of days using Canonical Correlation Analysis (CCA) ^20^. A Linear Discriminant Analysis (LDA) was used to classify the target locations (Fig. 1B). An LDA model was trained using data from Day N and tested within day (N on N) using leave one out cross validation (LOOCV). This analysis was then repeated for every possible pair of training day N and testing day M. Further materials and methods information can be found in the ‘Methods’ section of this manuscript.

**Figure 1.**
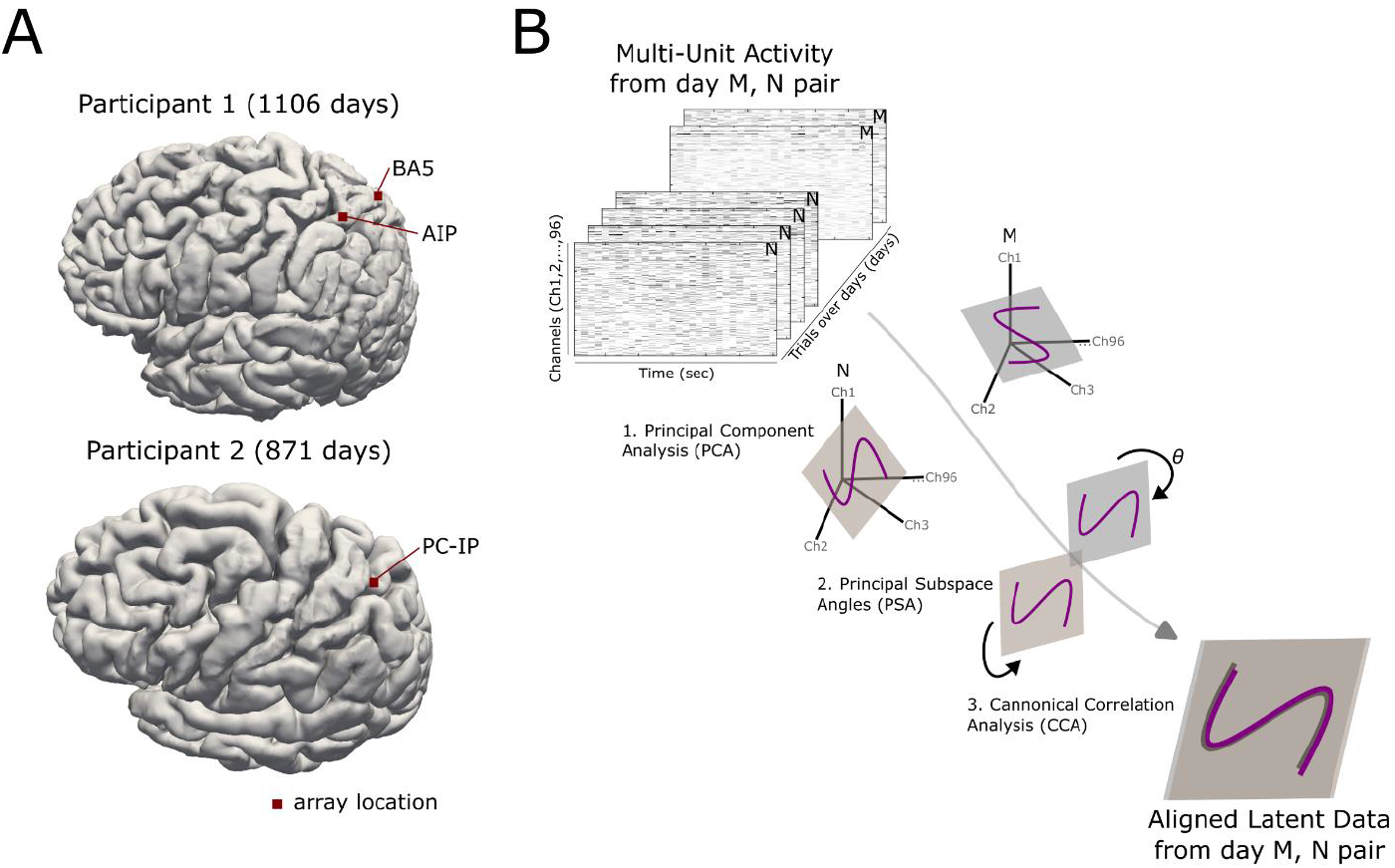
Methods. A) The neural signal was recorded on all days of the task using the same microelectrode arrays (Blackrock Neurotech). Arrays were implanted in the Anterior Intraparietal Area (AIP) and Brodmann Area 5 (BA5) of participant 1, and the junction of the postcentral and intraparietal sulcus (PC-IP) in participant 2. B) Data were arranged in a tensor of multi-unit activity (MUA) on each channel, of each trial completed during the duration of the study. All trials within a day were grouped and a pair of days was selected for further analysis (e.g., day N or M). For each day, the latent neural data was calculated using principal component analysis (PCA). The latent data were aligned by canonical correlation analysis (CCA), and the magnitude of the alignment was calculated as the principal subspace angle (PSA).

When decoding the MUA signal, we observe good decoding accuracy (Fig. 2A, MUA – red) within the same day, but this accuracy quickly degrades as the number of days between training and testing day increases (Fig. 2A, MUA – black). Intriguingly, aligned latent neural activity space substantially improves the accuracy (Fig. 2A, Latent). In particular, across all pairs of days, the decoding performance that can be achieved is higher from the latent signal (mean±SD, AIP: 51.2±8.38%, BA5: 63.7±12.0% and PCIP: 45.8±8.63%) than that achieved with MUA (mean±SD, AIP: 35.4±11.1%, BA5: 34.6±12.1% and PCIP: 25.5±11.8%) (all differences p<0.001, Wilcoxon Sign Rank test, Bonferroni corrected) (Fig. 2B). Further, the across-day training produces a comparable performance compared to within-day using aligned latent data. Across all recording electrode arrays, the correlation between performance and time between the pairs is smaller for latent signals (AIP r = -0.066, BA5 r = 0.020, PCIP = -0.033, Pearson’s linear correlation coefficient) than for MUA (AIP r = -0.12, BA5 r = -0.30 PCIP r = -0.29, Pearson’s linear correlation coefficient). To summarize these results, we calculate the ratio of performance between all the within-day models and all the across-day models for latent and MUA activity. A ratio of 1 represents a comparable result, a value greater than one would mean that across-day pairs performed better than within-day pairs and vice versa for values below 1 (Fig. 2C). Here we see that in all cases the ratio of latent signals is higher than MUA demonstrating the aligned latent signal across days has significantly increased stability compared to the MUA activity (p<0.001, Wilcoxon signed rank test, Bonferroni corrected. Participant 1 N = 20449, Participant 2 N = 5329).

**Figure 2.**
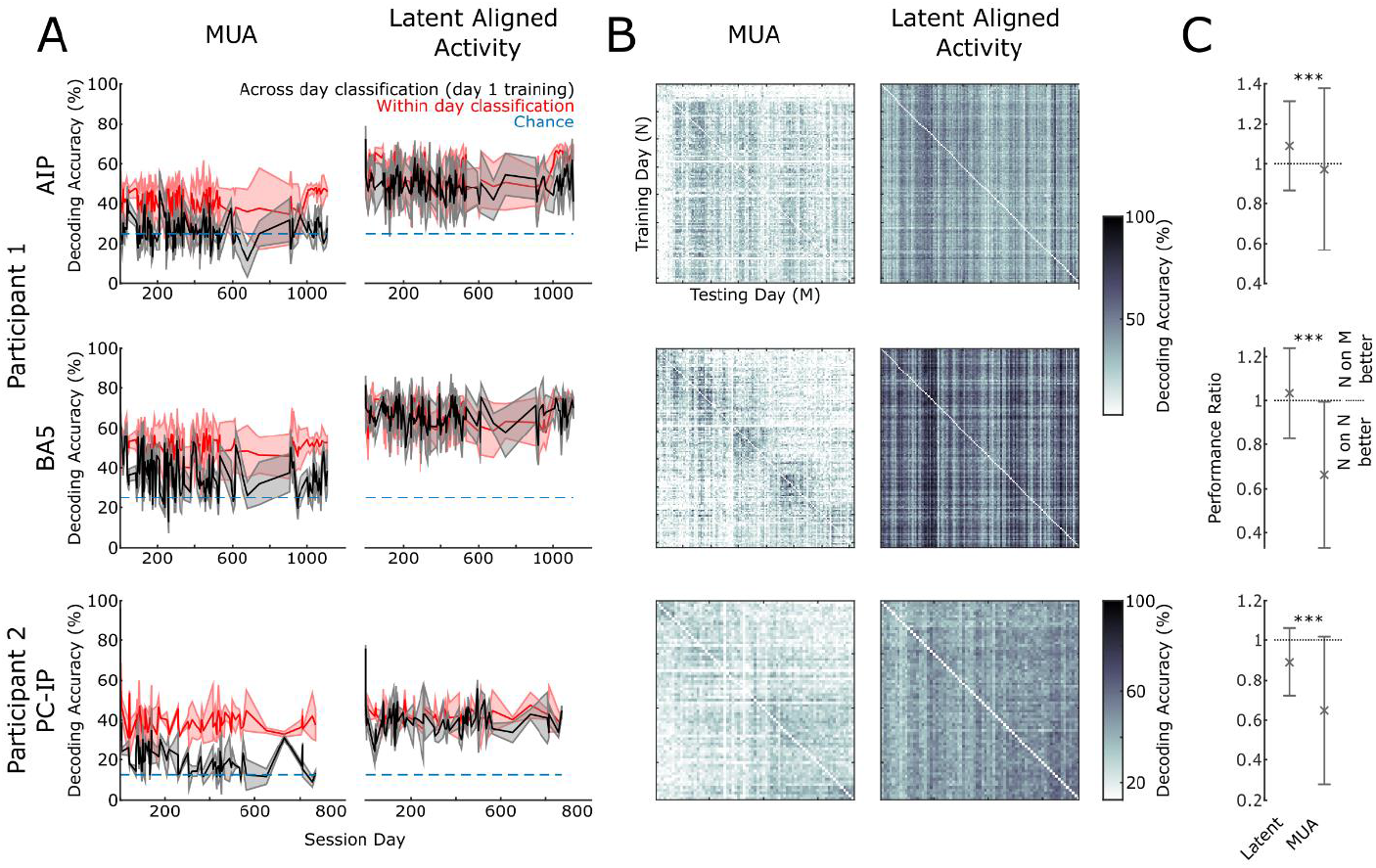
Classification decoder accuracy for two human participants. A) Left column: the performance of the LDA classifier using the raw multi-unit activity (MUA) from the brain regions AIP (top), BA5 and PCIP (bottom). The red line shows the classification accuracy when the data from the same day (N on N) is used for training and testing with leave-one-out cross validation. The black line shows the classification accuracy when data from session day N is used for testing, but training is always performed on data from a single day (in this example day 1). Shading shows the standard deviation, dotted line shows the chance level for classifications. Right column: the same analysis as the left column but using the latent aligned data. B) The decoding accuracy of every pair of days for MUA and Latent activity. C) The ratio of performance between within day and across day decoding, error bars show standard deviations, stars indicate significance.

In this task, the participants were not required to learn anything novel, so we do not expect neural activity changes related to learning. However, the repetitive nature of the task and the amount of time spent performing it may still alter the brain activity representing the task over time. To investigate the way in which the task is represented in the latent signal over time, we calculated the Principal Subspace Angle (PSA) ^21,22^ between all pairs of days (Fig. 3A). This angle reflects the magnitude to which the subspaces in the pair must be rotated to be maximally correlated, which is one way to measure the extent to which CCA rotates the latent neural data to align day pairs ^23^. A smaller PSA indicates a more similar representation between pairs. We computed the first PSA in participant 1 only, who had two arrays in different functional areas. We controlled for changes in the health of the array, which were not significantly different for the two arrays over time (see Methods Fig. 1C). We divided up the data into early and late periods (Methods Fig. 1A), where late was defined as the resumption of the experiment after a significant break (akin to a ‘washout’ in the familiarity of the task). Over all pairs of days, the variability in PSA in BA5 was smaller than for AIP (Fig. 3B left column). We then focused on close pairs of days within the early and late phases of data collection, those with a difference of only up to 10 days (Fig. 3B right column). Interestingly, after the break in the experiment, we find a significant difference in the representation of the data across these relatively close day pairs in only area AIP and not area BA5 (p<0.001, Wilcoxon Signed Rank Test). This finding indicates a more stable intrinsic representation of reaching in area BA5 than AIP.

**Figure 3.**
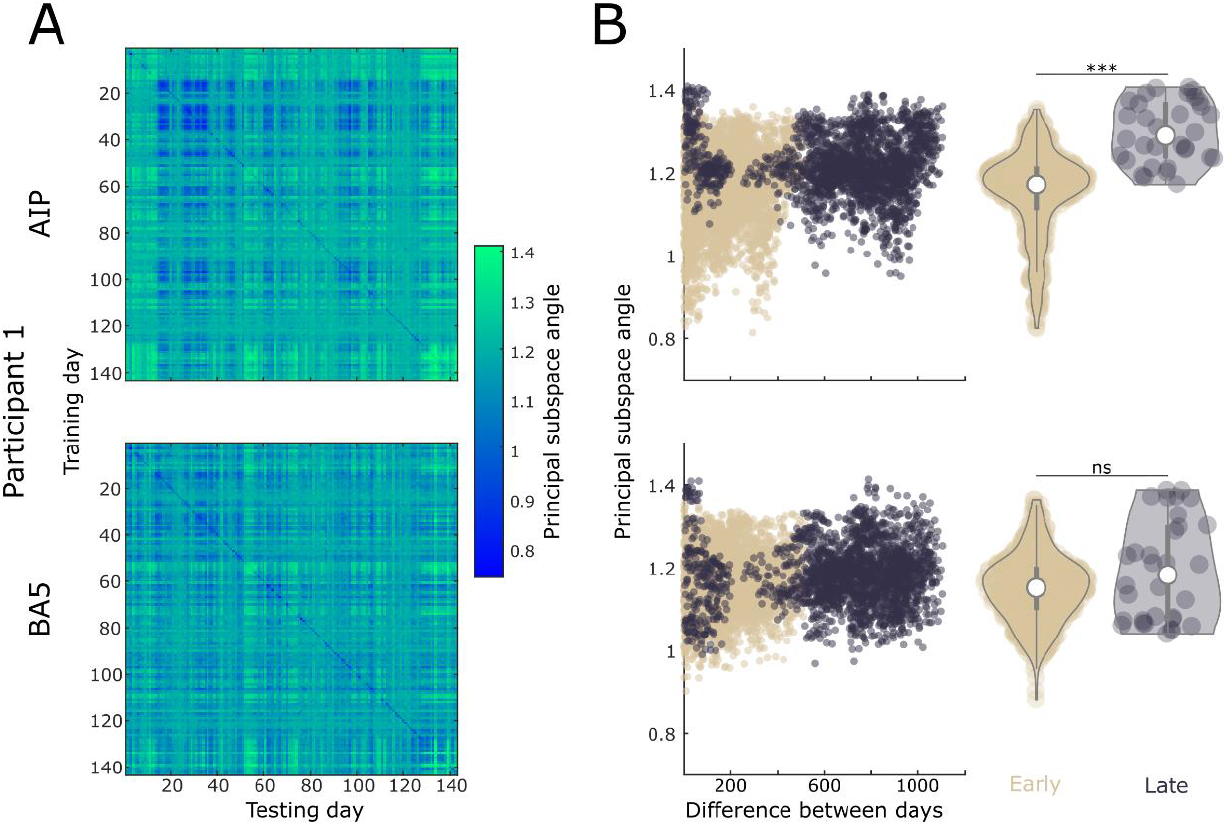
Principal subspace angles. A) The principal subspace angle (PSA) calculated between all pairs of days for AIP (top) and BA5 (bottom). B) Left Column: the PSA between each pair of days, colored according to the early and late period (Methods Fig 1A.) Right Column: violin plots of only the day pairs where the difference between days is 10 or less, grouped into the early and late period.

Here we have demonstrated the stable representation of neural activity in subspaces of human intracortical recordings over several years. This result validates methods of aligning latent spaces developed in non-human primates during actual reaching ^12,24,25^, here applied successfully in imagined reaches by humans. Furthermore, we have extended the finding of stable subspaces beyond the primary sensorimotor cortices into higher-order association areas in humans. The aligned latent signal performs best in decoding overall in each site, but the magnitude of the improvement reduces as the recordings come from more cognitive brain regions. We see the same effect in the PSA, where the variability in representation increased in the more cognitive brain region. We hypothesize that this trend is due to an increased flexibility in neuronal processing facilitating higher order/conceptual aspects of reaches (in AIP) compared to a more fundamentally engrained processing in the lower order sensorimotor control of a limb (in BA5).

There are few human intracortical BMI datasets available to validate the performance of latent signals over substantial periods of time in the same task. Participants enrolled in intracortical clinical trials are typically encouraged to perform a much broader range of tasks, with each requiring little to no training. Consequently, far fewer trials are available for any specific experimental paradigm. Chronic experiments exploring the human cortex offer a unique opportunity to study various changes in neural circuits over extended periods of time. Based on our results, we encourage the design of future studies to facilitate longitudinal task data collection and data collection from cortical sites beyond the traditionally used primary motor and sensory cortices. As we show here, these data allow the identification and validation of stable neural features which enable the long-term use of BMIs without the need for retraining, thus paving the way for BMIs to be used in many more cases; by individuals who lose the ability to retrain due to degenerative condition, or those who suffer injuries that preclude electrode implants in primary sensorimotor cortex. With the identification of such robust features, one promising direction for future work may be to enable the development of generalized BMIs that can be trained on data from individuals other than the eventual intended user ^26^.

## Methods

All procedures were approved by the Internal Review Boards of California Institute of Technology, University of Southern California, Rancho Los Amigos National Rehabilitation Center, University of California Los Angeles and Casa Colina Hospital and Centers for Healthcare. Informed consent was obtained from all participants after the nature of the study and possible risks were explained. This work was performed as part of Clinical Trials: NCT01849822, NCT01958086, NCT01964261.

### Participants

Participant 1 was a 32-year-old tetraplegic male at the time of implantation. He was implanted with two microelectrode arrays on 17 April 2013. The electrodes were implanted in Brodmann Area 5 (BA5) and the Anterior Intraparietal Area (AIP). He had a complete lesion of the spinal cord at cervical level C3-4, sustained 10 years earlier, with paralysis of all limbs. Participant 2 was a 59-year-old tetraplegic female at the time of implantation. She was implanted with two arrays but only one was used in this study, at the junction of the post-central and intraparietal gyrus (PC-IP) on 29 August 2014. The other array was not functional. She had a C3-C4 spinal lesion (motor complete) sustained 7 years earlier, and retained movement and sensation in her upper trapezius, without control or sensation in her hands. During their enrollment, the participants performed many different tasks. The data for the analysis presented in this manuscript were collected on 143 and 73 unique days, over a period of 1106 and 871 days, for participant 1 and 2, respectively.

### Task and Data Collection

The center out task was intended to allow the participants to spatially position a cursor on a computer screen. Targets were presented one at a time on the LCD display. The LCD monitor was positioned approximately 184cm from the subject’s eyes. Stimulus presentation was controlled using the Psychophysics Toolbox for MATLAB. During recording-only sessions, without any decoder, a circular cursor on the screen would move automatically from the center to one of either 4 (participant 1) or 8 (participant 2) targets arranged radially around the center point. Following a 250ms delay relative to target onset, the cursor moved in a straight line directly to the target with an approximately bell-shaped velocity profile. Each trial lasted 3 seconds. The number of trials completed by the participants on each day during the study is shown in Methods Fig. 1A and B. Participants were asked to imagine making movements of the arm to mimic the movements observed on the screen.

The NeuroPort System (Blackrock Neurotech, UT, USA), comprising the arrays and neural signal processor (NSP), has received Food and Drug Administration (FDA) clearance for <30 days of acute recordings. For this study we received FDA IDE (Investigational Device Exemption) clearance for extending the duration of the implant. The health and performance of the arrays was assessed as the mean impedance across all electrodes on each array, recorded on each day of the experiment.

Impedance data is available for participant 1 only (Methods Fig. 1C). Multi-Unit Neural Activity (MUA) was amplified, digitized, and recorded at 30 kHz with the NeuroPort NSP. The threshold for calculation of MUA spikes was -3.5 * root mean squared voltage, calculated over each recording session. Data was organized into a three-dimensional tensor; The first dimension was the MUA binned into non-overlapping 50ms windows. The second dimension was the number of electrodes (96 for each array). The third dimension was the index of trials ordered chronologically from the first to last over the entire study period.

### Analysis

The analysis methods used in this manuscript extend the analysis of Gallego and colleagues ^12^ who demonstrated success in long-term decoding from primate motor cortex recordings. Data from each participant and each array was processed separately. The analysis was completed identically between all pairs of all days in which the participants completed the center out task. For the following description of the analysis two such days are represented as day M and day N.

Initially the same number of trials, containing equal presentation of all targets, are taken on each day. On days with different numbers of trials, we used all the trials on the day with the fewer trials and then randomly selected the same number of trials from the other day. To ensure all the trials for a pair of days were used, the entire analysis was repeated 1000 times, each iteration using a different randomly selected set of trials. All electrodes (96) and all time bins were included for all trials. For each day this produced a (electrode x time x trials) matrix. The data was concatenated across trials and then dimensionality reduction was performed using Principal Components Analysis (PCA) (‘pca’ function, Matlab 2021b). We reduced the data to 10 dimensions, following previous analysis, but confirmed that the results did not qualitatively change using a larger range of values. The result of the PCA analysis was a (10 x time*trials) matrix. We call this the ‘latent data’. The latent data from each day in the pair was then aligned using Canonical Correlations Analysis (CCA) (‘canoncorr’ function, Matlab 2021b). This produced a (10 x time*trials) matrix. We call this the ‘aligned latent data’. The data was then split back into individual trials (10 x time x trials) and the activity in each trial (‘time’ dimension) was averaged producing a (10 x 1 x trials) matrix for each day M and N. This was then used to calculate a linear regression model for classification (‘fitlm’ function, Matlab 2021b). The aligned latent data were used as the data and the target labels for each trial were used as the model. For within day calculations of classification accuracy a leave-one-out cross validation (LOOCV) was used to calculate classification accuracy. For calculating classification accuracy across days, the entire data from day M was used to train the LDA model, which was tested on the entire dataset from day N (and vice versa). To calculate the Principal Subspace Angle (PSA) we followed the method presented by Knyazev and colleagues ^21^ (‘subspacea’ function, MATLAB Central File Exchange, Matlab 2021b). We present data from the first PSA, but we note the results remained qualitatively consistent when summing over all PSAs calculated from the data.

We only perform the PSA analysis on participant 1 because two electrode locations were recorded. In this case AIP and BA5 control for each other in factors related to changes in electrode-tissue interface that could influence the results (see Methods Figure 1C). We assume that since this crucial metric is consistent between the two arrays analytical differences can be explained by the different neurophysiology of the regions. Since there is only the PCIP array in participant 2 we cannot make the same assumption about differences over time, as without a comparison, analytical differences may be due to changes in the electrode-tissue interface.

## Data availability

The datasets analyzed for this manuscript will be shared upon reasonable request.

## Code availability

All analyses were implemented using custom Matlab (The Mathworks Inc.) code. Code to replicate the main results will be shared upon reasonable request.

---

**Methods Figure 1.**
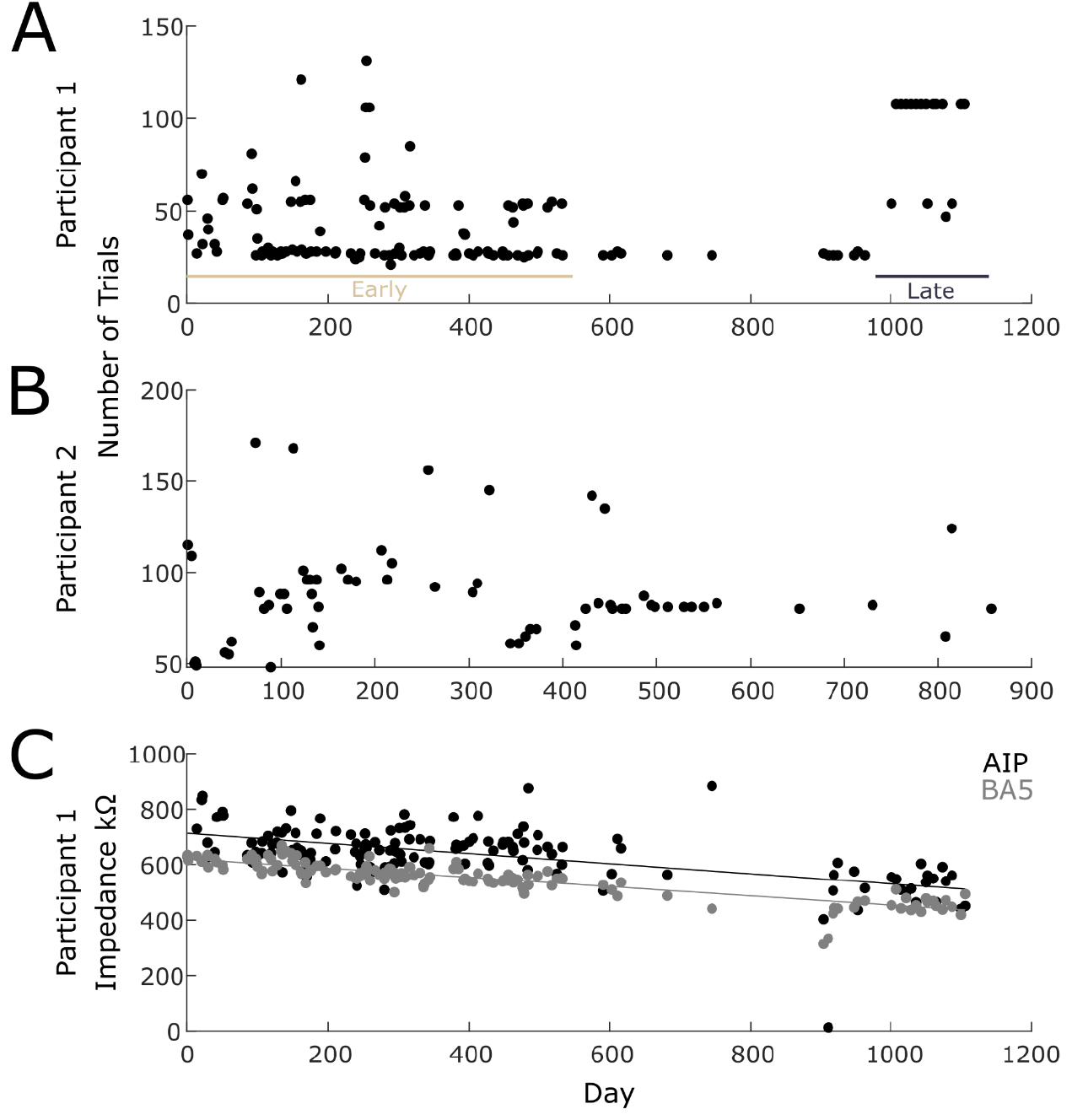
Data collection. A & B) Varying numbers of trials of the same task were collected on each day. Participant 1 only – during a middle period relatively few trials of the task were performed to focus on other experiments. Data collected were split into an ‘Early’ and ‘Late’ period. C) The impedance of each array on each day experiments were collected. Impedance data are only available for participant 1.

## Acknowledgements

The authors would like to acknowledge the outstanding participation of NS and EG, whose involvement in the study over many years provided these data sets. We would like to acknowledge the neurosurgical teams responsible for the implants: Participant 1; Brian Lee (USC) and Charles Liu (USC) and Participant 2; Nader Pouratian (UCLA). We would additionally like to thank all members and collaborators of the Andersen Lab who were involved in the collection of data during the enrollment of these participants (2013-2019). This work was supported by the National Institute of Health (R01EY013337, R01EY015545), The Boswell Foundation and the T&C Chen Brain-Machine Interface Center at Caltech.

